# Cortical control of locomotor muscle activity through muscle synergies in humans: a neural decoding study

**DOI:** 10.1101/413567

**Authors:** Hikaru Yokoyama, Naotsugu Kaneko, Tetsuya Ogawa, Noritaka Kawashima, Katsumi Watanabe, Kimitaka Nakazawa

**Author notes:** Correspondence to: Kimitaka Nakazawa Department of Life Sciences, Graduate School of Arts and Sciences, The University of Tokyo, 3-8-1 Komaba, Meguro, Tokyo 153-8902, Japan.

## Abstract

Walking movements are orchestrated by the activation of a large number of muscles. The control of numerous muscles during walking is believed to be simplified by flexible activation of groups of muscles called muscle synergies. Although significant corticomuscular connectivity during walking has been reported, the level at which the cortex controls locomotor muscle activity (i.e., muscle synergy or individual muscle level) remains unclear. Here, we examined cortical involvement in muscle control during walking by brain decoding of the activation of muscle synergies and individual muscles from electroencephalographic (EEG) signals using linear decoder models. First, we demonstrated that activation of locomotor muscle synergies was decoded from slow cortical waves with significant accuracy. In addition, we found that decoding accuracy for muscle synergy activation was greater than that for individual muscle activation and that decoding of individual muscle activation was based on muscle synergy-related cortical information. Taken together, these results provide indirect evidence that the cerebral cortex hierarchically controls multiple muscles through a few muscle synergies during walking. Our findings extend the current understanding of the role of the cortex in muscular control during walking and could accelerate the development of effective brain-machine interfaces for people with locomotor disabilities.

## Introduction

Human locomotor movement is organized by the coordinated activation of a large number of muscles. It has been suggested that complex muscle activity is generated from a small number of groups of muscle activations called muscle synergies [1-5]. Locomotor muscle synergies are thought to be structured in the spinal circuitry [6, 7]. Based on previous studies examining synergy activation among different subject groups, it has been suggested that the cortex activates locomotor muscle synergies [1, 2, 7]. These studies reported that locomotor muscle synergy in healthy adults exhibited activation that was sharply timed around gait events [1], whereas locomotor muscle synergy in neonates [2] and complete spinal cord injury (SCI) patients [7] exhibited smooth prolonged activation. The differences in the patterns in neonates and SCI patients could be caused by immature and injured corticospinal pathways, respectively. Based on these findings, it is thought that cortical descending commands modulate basic locomotor muscle synergy activation generated by subcortical structures, particularly in the spinal cord. However, there is currently no direct evidence of cortico-muscle synergy relationships supported by simultaneous recordings of cortical activity and muscle synergy activation during walking.

Unlike quadruped animals [8, 9], human bipedal walking is characterized by significant cortical activity even during undemanding steady-state walking [10-19]. Significant cortical activation has been demonstrated previously in premotor, supplementary motor, and primary sensorimotor regions during real and imagined walking using neuroimaging techniques such as positron emission tomography (PET) and near-infrared spectroscopy (NIRS) [11, 12]. Recent studies using electroencephalography (EEG), which has greater temporal resolution, have demonstrated gait-phase dependent modulation of cortical activity, particularly in the sensorimotor cortex, using a combined method of independent component analysis and source localization techniques [13-17]. Other EEG studies have demonstrated significant corticomuscular connectivity between the leg sensorimotor area and leg muscles during walking using individual muscle level analysis [18, 19]. Although these studies strongly suggest cortical involvement in muscular control during walking, at what level the cortex controls muscle activity remains unclear, i.e., at muscle synergies or individual muscles.

To address the question, we hypothesized that the human cortex controls locomotor muscle activity through muscle synergies, rather than through direct control of each muscle and, then, examined how the cortex is involved in muscle control during walking by decoding the activations of muscle synergies and individual muscles from EEG signals. Brain decoding techniques, which predict the mental or motor state of a human from recorded brain signals, have received substantial attention for the development of brain-machine interfaces (BMIs) for repairing or assisting deficits in cognitive or sensory-motor functions [20-22]. In addition to potentially restoring lost functions, neural decoding can provide information on the physiological principles of how motor movements are controlled by the brain [23].

In this study, using neural decoding techniques, we demonstrate that the activation of muscle synergies can be decoded from cortical activity and that the decoding accuracy for muscle synergies is greater than that for individual muscles. Additionally, we show the decoding of individual muscle activity is based on muscle synergy related cortical information. These results provide experimental evidence that the cortex hierarchically controls individual muscles through locomotor muscle synergies. The findings will shed light on the cortex’s role in muscular control during walking and provide an important basis for developing effective neuroprostheses for walking rehabilitation.

## Results

Healthy participants walked on a treadmill at 0.55 m/s for 7 min 30 seconds. Surface electromyographic (EMG) signals were recorded from 13 leg muscles on the right side. EEG signals were recorded from 63 channels. EEG data from 30 channels (Figure 1), which are assumed to be less affected by eye blinks and facial/cranial muscle activity, were used for subsequent analysis. Using the EMG and EEG signals, we tried to decode individual muscle and muscle synergy activations from cortical activity. See Figure 2 for an overview of our decoding methodology.

**Figure 1.**
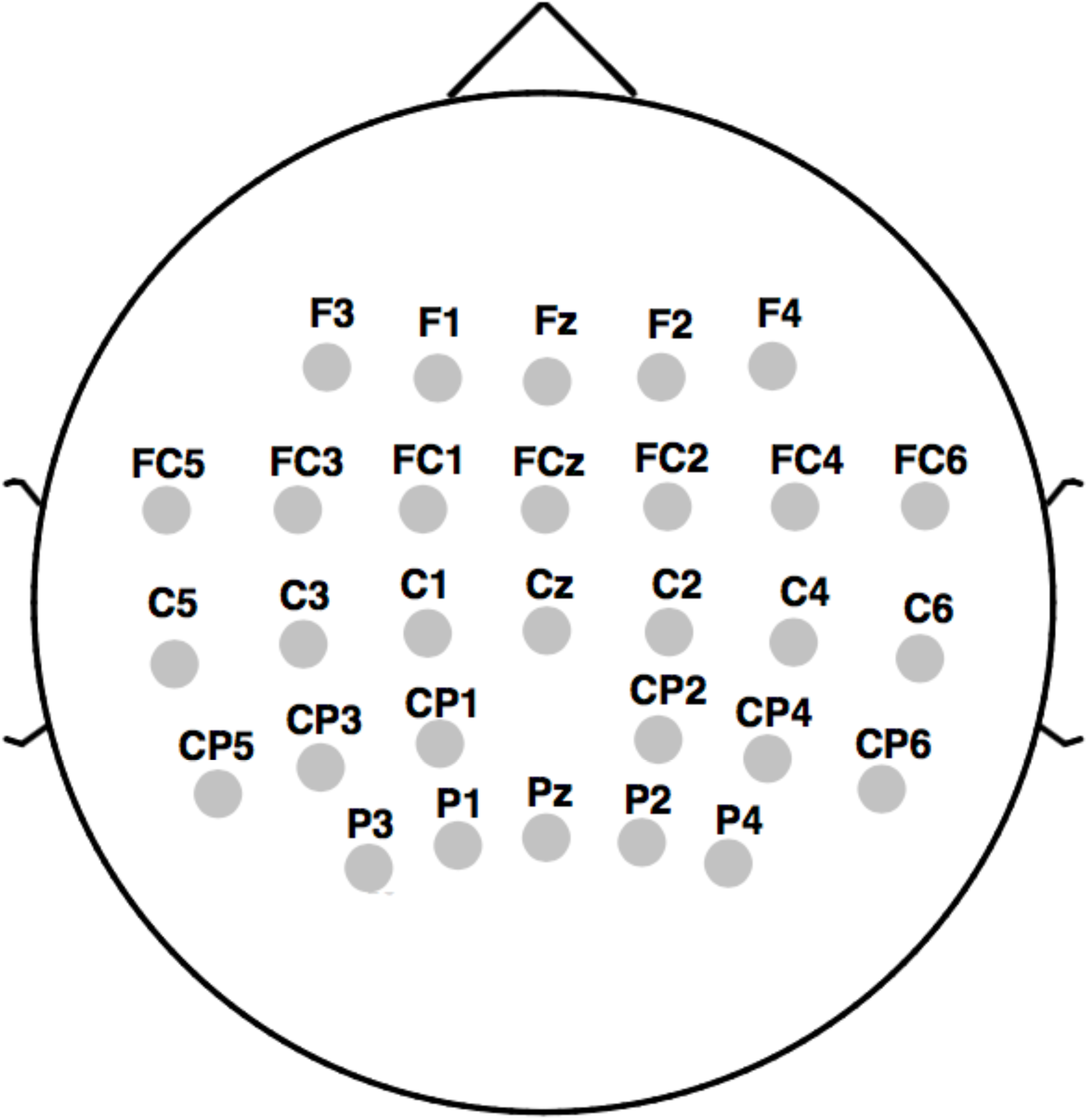
EEG electrode montage corresponding to the international 10-20 system.

**Figure 2.**
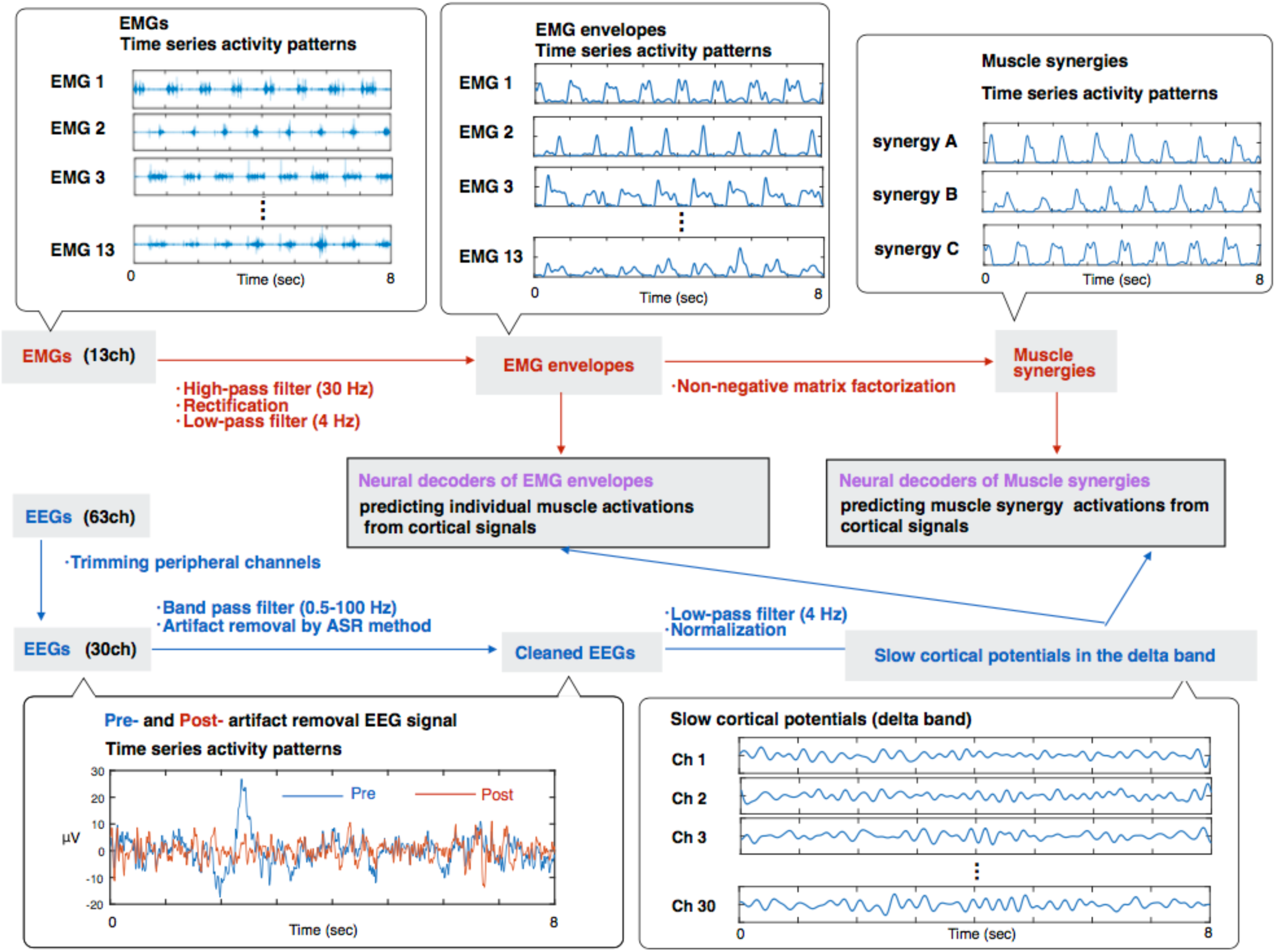
Schematic diagram depicting the neural decoding of locomotor muscle synergy and individual muscle activations from simultaneously recorded EEG signals. Examples of 8 seconds of raw EMG signals, EMG envelopes, muscle synergies, pre- and post-artifact removal EEG signal from an electrode, and slow cortical potentials in the delta band are shown.

### Extracted locomotor muscle synergies

The recorded EMGs were rectified and smoothed by a low-pass filter. Next, using non-negative matrix factorization (NMF) [2, 3, 24-26], muscle synergies were extracted from each participant. From the low-pass filtered EMGs, 4.17 ± 0.58 (mean ± SD) muscle synergies were extracted from each participant. The extracted muscle synergies were grouped into five types using cluster analysis (synergy A–E, Figure 3). Supplemental Table S1 summarizes the characteristics of the extracted locomotor muscle synergies.

**Figure 3.**
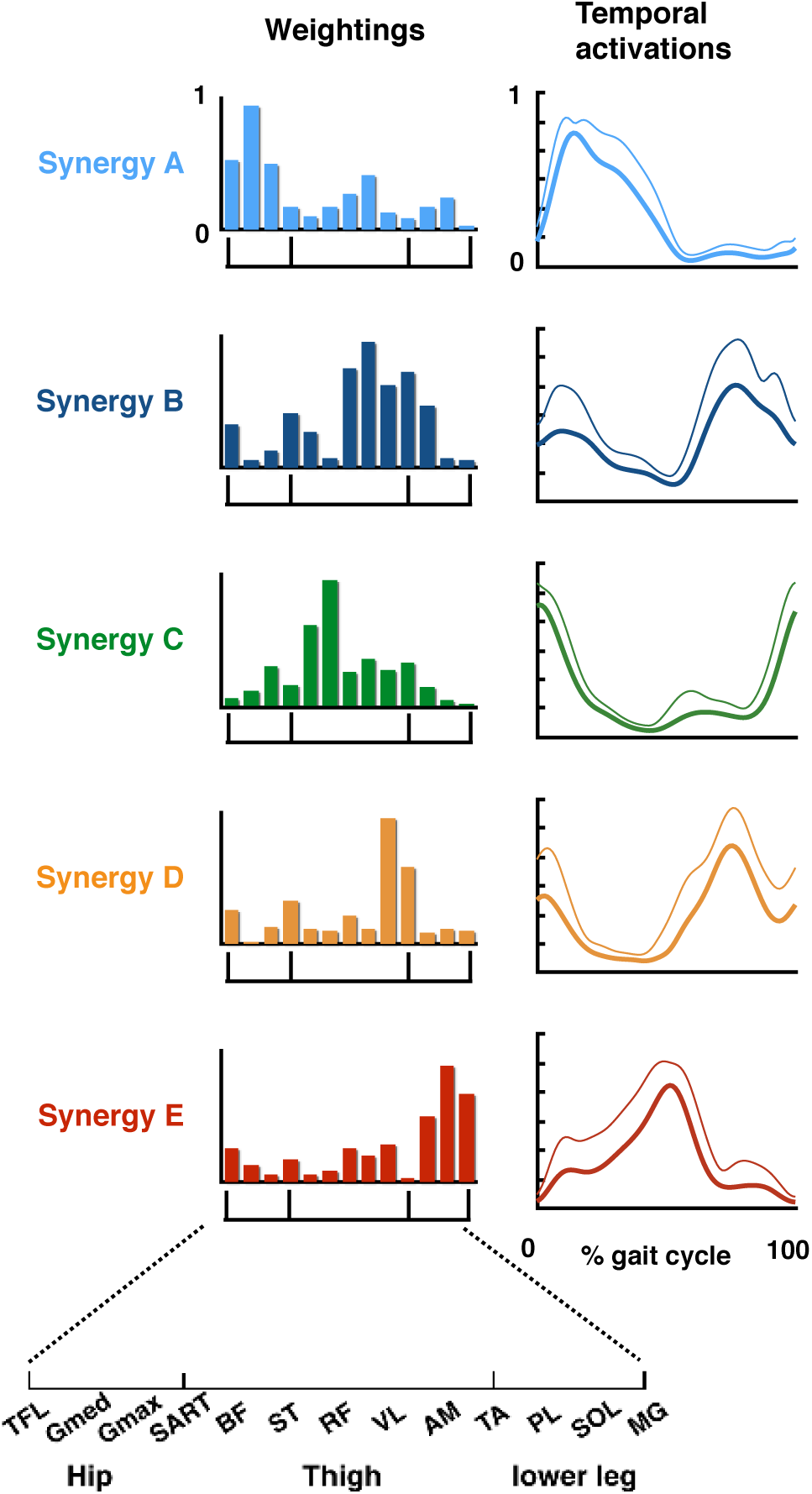
Five extracted types of locomotor muscle synergies. Average muscle weightings (bars) and corresponding temporal activation patterns (waveforms) across participants in each type of locomotor muscle synergy are shown. Each bar height represents the relative level of activation of each muscle synergy. An enlarged view of the x-axis is shown at the bottom. Lines indicate the temporal activation patterns of the muscle synergies. Thick lines indicate average temporal activation patterns, while thin lines indicate their standard deviations (SD).

### Neural decoding of activation of muscle synergies and individual muscles from EEG signals

As preparation for neural decoding, recorded EEG signals were band-pass filtered in the delta bend (0.5-4Hz). The filtered signals, which are called slow cortical potentials, were confirmed to be particularly informative for decoding motor-related parameters [27-32]. We used multiple linear models, also referred to as Wiener filter, to decode individual muscle and muscle synergy activations from the slow cortical potentials, as used in previous studies decoding motor parameters [27-32]. Figure 4 provides examples of real and reconstructed muscle synergy activations (Figure 4A) and individual muscle activations (Figure 4B) from a participant. In this participant, all locomotor muscle synergy activations were successfully reconstructed based on visual inspection (Figure 4A). In contrast, in individual muscle activation, the amplitude modulation was not sufficiently reconstructed in some muscles, such as SART, AM, PL, SOL (Figure 4B).

**Figure 4.**
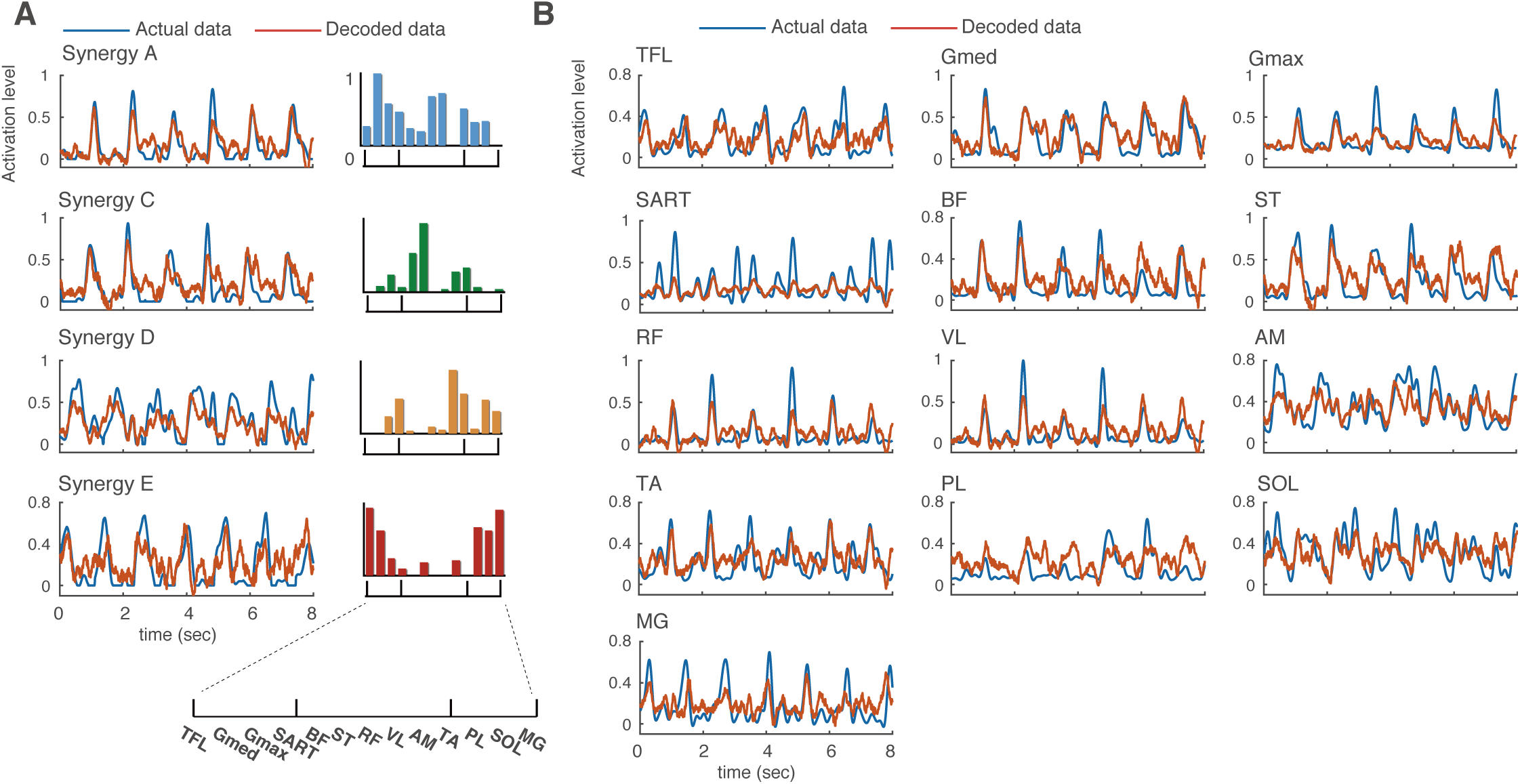
Typical examples of decoded and actual muscle synergy activations (A) and individual muscle activations (B) from a participant. Red and blue waveforms indicate decoded and actual activation patterns, respectively. Bars represent muscle synergy.

To quantify decoding accuracy, we calculated the Pearson’s correlation coefficients (*r*) between the real and reconstructed activations in each decoder (Figure 5). The mean values across the participants ranged from 0.48 to 0.52 in muscle synergy decoders and 0.31 to 0.52 in individual muscle decoders (Figure 5A). The overall accuracy (i.e., averaged correlation values across all decoders in each type [muscle synergy or individual muscle]) of the muscle synergy decoder was higher than that of the individual muscle decoder (*t*(11) = 5.30, *p* = 0.0003, paired t-test, Figure 5B).

**Figure 5.**
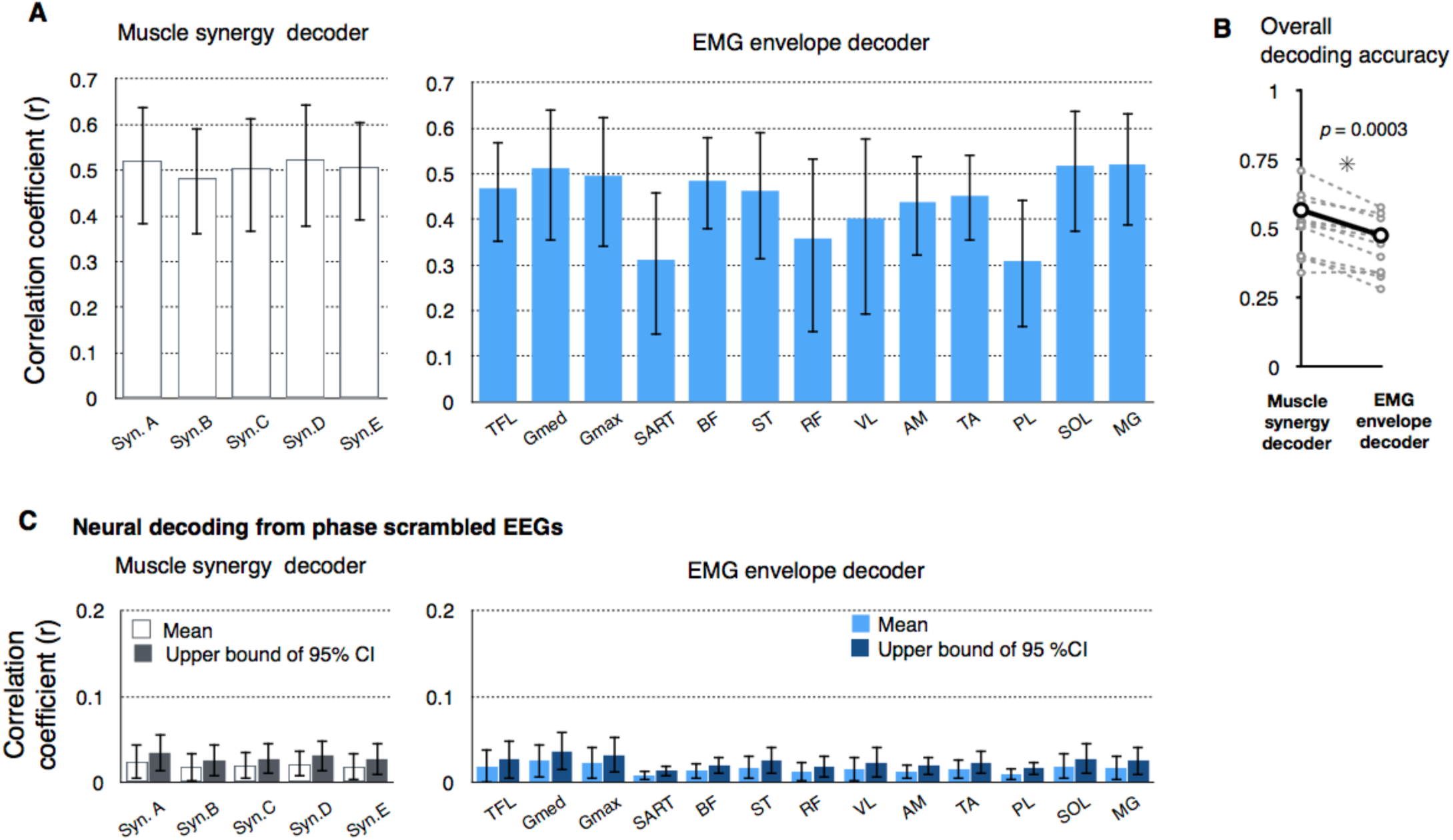
Decoding accuracy of activation of muscle synergies and individual muscles. (A) Decoding accuracy (correlation coefficient) for each muscle synergy type (left) and EMG envelope of an individual muscle (right). The mean and SD across participants are shown. (B) Overall decoding accuracy for muscle synergy decoders and individual muscle decoders. Mean values across participants (black) and each participants’ data (gray) are shown. (C) Decoding accuracy when EEG phase was scrambled. The bars indicate the participant’s mean of means and the upper ends of the 95% confidence interval obtained from the distribution of the surrogate datasets of EEG signals.

Next, to validate the results of neural decoding, the same decoding process was performed on phase-randomized EEG signals to estimate the chance levels. We generated 100 surrogate datasets and evaluated the mean and 95% confidence intervals of the decoding accuracy from the distribution of decoding accuracy of the surrogate datasets (Figure 5C). The decoding accuracy from the phase-randomized data was low regardless of the type of muscle synergies or individual muscles (range of mean *r* values: 0.0085-0.025). The decoding accuracy from the original EEGs exceeded the 95% confidence interval of the surrogate datasets for all muscle synergy and individual muscle decoders in all the participants.

### Relationships between muscle synergy decoders and individual muscle decoders

Although the decoding accuracy of muscle synergy activation was similar for all synergy types, the decoding accuracy of individual muscle decoders varied widely across different muscles (Figure 5A). In this study, the cortex is assumed to be involved in muscle control through muscle synergies. Based on this assumption, the variability of decoding accuracy in individual muscles would be reproduced by individual muscle activations indirectly decoded from muscle synergy activations decoded from muscle synergy decoders. To test this hypothesis, we reconstructed individual muscle activations by summing the outputs of each decoded muscle synergy (Figure 6A). The decoding accuracy of directly decoded individual muscle activations was found to have a very strong positive correlation with that indirectly decoded from the outputs of decoded muscle synergies (*r* = 0.97, Figure 6B). This result indicates that if muscle activation is not well decoded through decoded muscle synergies, the decoding accuracy of the muscle will be low even when it is directly decoded.

**Figure 6.**
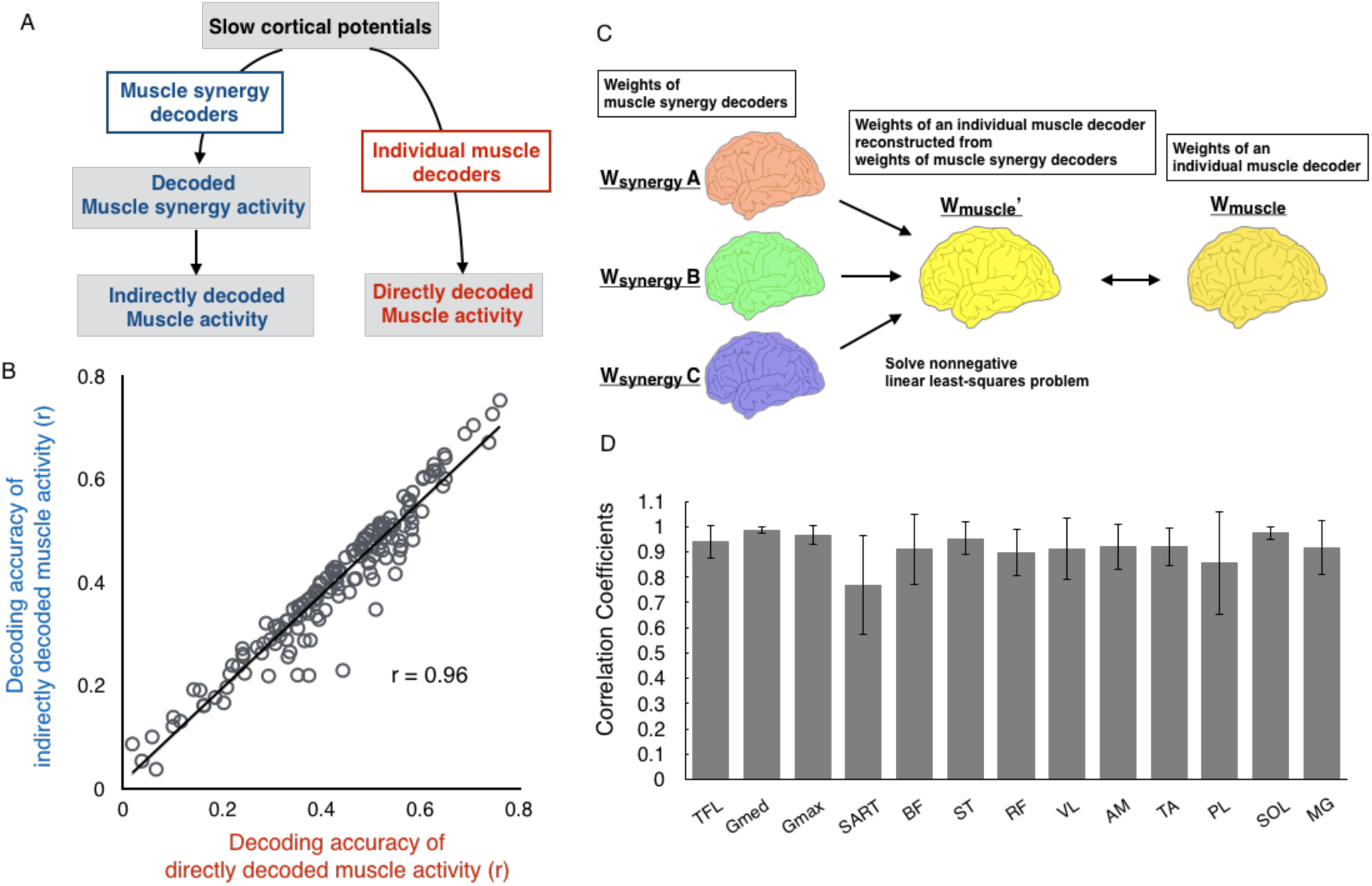
Relationships between muscle synergy decoders and individual muscle decoders. (A) A schematic flow diagram illustrating directly decoded muscle activity and indirectly decoded muscle activity reconstructed from decoded muscle synergies. (B) Relationships of decoding accuracy in each individual muscle activity between those directly decoded from individual muscle decoders and those indirectly decoded from muscle synergy activations decoded from muscle synergy decoders. Each plot indicates the value of an individual muscle from a participant. (C) Schematic diagram of reconstruction of weights of an individual muscle decoder from those of muscle synergy decoders. (D) Similarity of weights of individual muscle decoders between the originals and those reconstructed from weights of muscle synergy decoders. The mean and SD across participants are shown.

The decoding accuracy relationships suggest that decoding of individual muscle activation is based on muscle synergy-related cortical information. If so, the weights of the individual muscle decoders (*W*_*muscle*_) should be represented as a linear combination of those of muscle synergy decoders (*W*_*syn*_) with non-negative coefficients. To test this possibility, 300-dimensional weights of an individual muscle decoder were reconstructed as a linear combination of the weights of muscle synergy decoders with non-negative coefficients (*W*_*muscle*_*’*, conceptual schema presented in Figure 6C). The similarity between the original and reconstructed weights (i.e., *W*_*muscle*_ and *W*_*muscle*_*’*, respectively) was quantified by Pearson’s correlation coefficient, which was 0.91 ± 0.11 (mean ± SD) across all muscles of all the participants. Regarding each type of muscle, the mean similarity values across participants ranged from 0.77 to 0.99 (Figure 6D). Thus, as expected, the weights of individual muscle decoders represented very similar patterns as those reconstructed from the weights of muscle synergy decoders.

### Contributions of electrodes to neural decoding

To evaluate the spatial contributions of cortical activity for predicting muscle synergy activations, we calculated the contribution of each electrode from the weights of the decoding model [33]. Figure 7A shows examples of the contributions of each electrode to the decoding in one participant. In this participant, the contribution of each electrode was approximately 7% at the highest. Thus, widely-distributed cortical activity, rather than activity from one specific electrode and area, contributed to decoding. The widely-distributed contribution of the whole cortex was also observed in the mean contribution in each type of synergy (Figure 7B).

**Figure 7.**
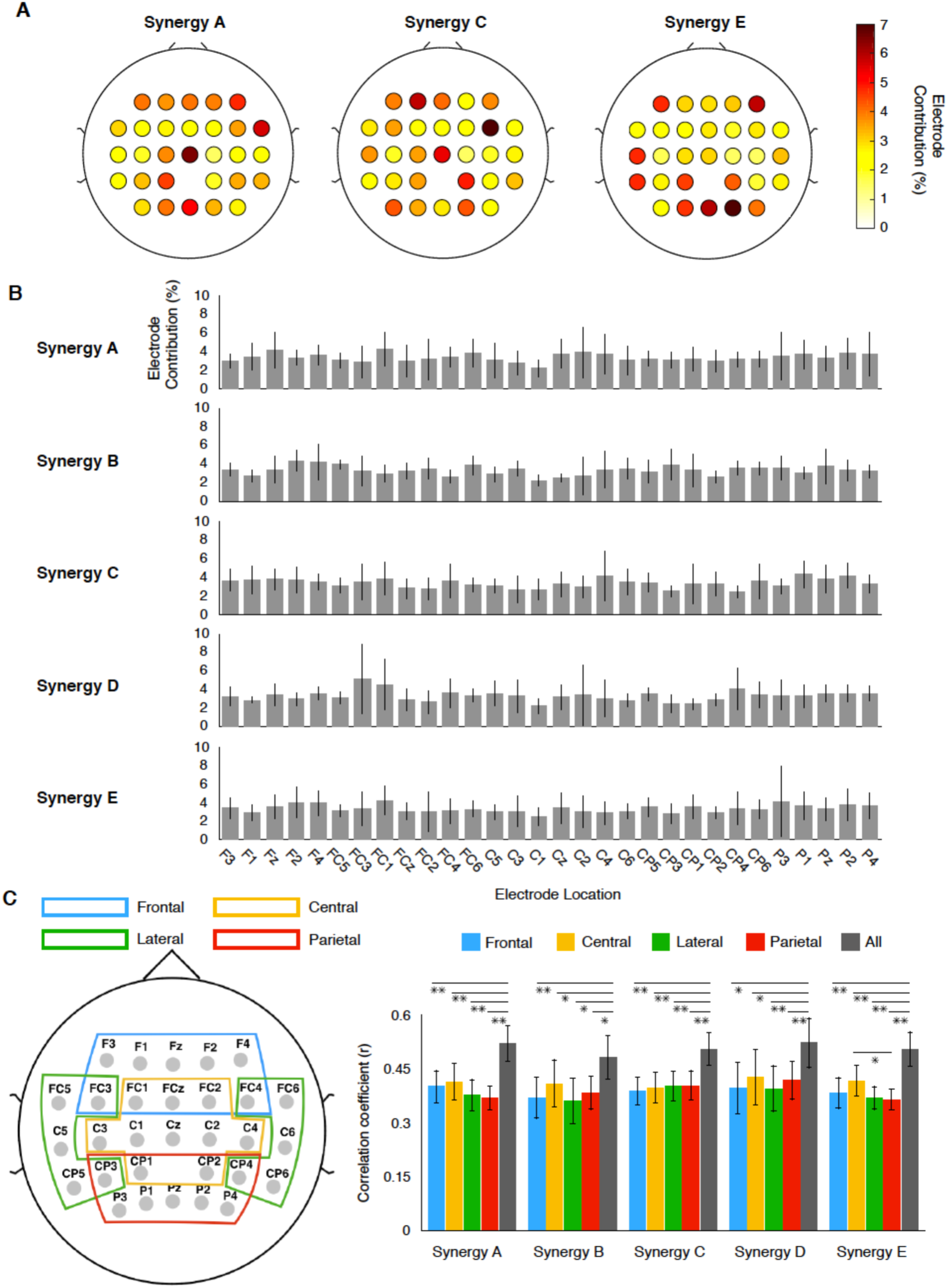
Contribution of each electrode to the decoding of muscle synergy. (A) Examples of contributions of each electrode to decoding from a participant (B) Mean contribution of each electrode in each synergy type. The error bars indicate the SD (C) Scalp map indicating the electrodes included in each region of interest (ROI) to examine the contributions from different cortical regions to decoding. (D) Decoding accuracy by each ROI and all electrodes. Data are represented as mean ± SEM. Asterisks indicate significant differences (*: *p* < 0.05, **: *p* < 0.01, FDR corrected for multiple comparisons, See also Table S2-S6 for detailed statistical values).

These results suggest that broad cortical activity is involved in the control of locomotor muscle synergy. To further validate the widely-distributed contribution of cortical activity to the decoding of locomotor muscle synergy, we divided the electrodes into four major regions of interest (ROIs), namely frontal, central, lateral, and parietal ROIs (Figure 7C). Next, for each ROI, we performed the same decoding procedure used for all electrodes and compared the decoding accuracies. The comparisons of the decoding accuracy did not show any significant differences among ROIs, except for that between the central and parietal ROIs in synergy E (Figure 7D). Nevertheless, the decoding accuracy in the full electrodes was significantly higher than that in each ROI (Figure 7D, *p* < 0.05, False discovery rate (FDR) corrected for multiple comparisons, see Table S2-S6 for detailed statistical values). Interestingly, the mean decoding accuracy of the central ROI was the largest for all synergy types except synergy C, and a significantly higher accuracy was found in the central ROI compared to the parietal ROI for synergy E (Figure 7D, *p* = 0.020, FDR corrected for multiple comparisons).

## Discussion

### Cortical control of locomotor muscle synergy

The last 15 years of research has suggested that cortical descending commands modulate basic locomotor muscle synergy activation generated by subcortical structures [1, 2, 7, 25]. Nevertheless, currently, there has been no evidence of cortical control of locomotor muscle synergies from simultaneously recorded cortical and muscle activity. In this study, we revealed that activation of locomotor muscle synergies decoded from EEGs was moderately correlated with real activation (Figure 5A), and that decoding accuracy of muscle synergy activation was generally higher than that of individual muscle activation (Figure 5B). By examining the relationships between individual muscle and muscle synergy decoders, we also showed that the decoding of individual muscle activity is based on muscle synergy-related cortical information (Figures 6B and 6D). Combined, the decoding results demonstrate significant cortico-muscle synergy relationships during walking, thus supporting the hypothesis that the human cortex hierarchically controls locomotor muscle activity through muscle synergies rather than by directly controlling each muscle [34].

Regarding cortical control of locomotor muscle synergies in a stroke-injured neuromuscular system, fewer locomotor muscle synergies resulting from the merging of healthy muscle synergies are recruited, which reflect disruption in the corticospinal descending pathways [25]. Therefore, post-stroke changes suggest cortical involvement in the activation of the locomotor muscle synergies. Other evidence regarding the cortical control of locomotor muscle synergies has been suggested by altered activation of the muscle synergies in patients with complete SCI [7] and neonates [2]. Both subject groups exhibited smooth sinusoidal-like activation patterns of locomotor muscle synergies rather than sharply timed activation, which was observed in healthy subjects [1, 35]. The sinusoidal-like activation patterns were also observed in other mammals [2]. Based on luck of corticospinal interactions in SCI patients and neonates, this similarity may suggest that the sinusoidal-like activation patterns are phylogenetically conserved in the spinal circuits. Taken together, it is possible that cortical descending commands modulate basic locomotor muscle synergy activation patterns generated by the spinal cord into the sophisticated patterns underlying human-specific upright bipedal walking.

The decoded muscle synergy activation observed here exhibited moderate correlation with actual activation (Figure 5). This moderate decoding accuracy is expected to derive from muscle synergy recruitment via multiple neural pathways, such as the brainstem, spinal cord, and sensory feedback [34, 36, 37] in addition to the cortex. Although the cortex is likely to be involved in the control of locomotor muscle synergies, its contribution may not be exclusively dominant. Partial contribution of the cortex to the control of locomotor muscle synergies may explain the moderate decoding accuracy observed in this study.

### Global cortical involvement in control of locomotor muscle synergies

Widely-distributed cortical activity, rather than activity from a specific electrode or area, contributed to decoding (Figure 7). Similarly, previous studies of neural decoding while walking demonstrated that leg kinematics could be decoded from cortical signals from widely-distributed regions [27, 28]. Previous work has shown the contributions of widespread cortical circuits, including the posterior parietal cortex, motor cortex, somatosensory cortex, and visual cortex, to visually guided walking in cats [38]. Although the contribution of widespread cortical circuits is limited to challenging walking conditions in cats, such circuits may contribute to the control of human walking even during steady-state walking because the mechanical instability of human-specific bipedal walking [39] requires additional cortical involvement. Indeed, widespread cortical activity has been reported during human walking by source estimation of EEG signals [13, 40]. In addition, motor imagery studies have demonstrated locomotor-related activity in brain regions including the primary and supplementary motor cortex and several bilateral parietal and frontal regions using functional magnetic resonance imaging (fMRI) [41, 42]. Thus, it is possible that locomotor-related global activity in the cortex can explain the widely-distributed contribution of electrodes to the decoding of locomotor muscle synergy activations in the present study.

Of note, the mean decoding accuracy of the central ROI was highest among all ROIs in all synergy types except one, and significantly higher accuracy was found in the central ROI compared to the parietal ROI in one synergy type (Figure 7D). Since the central ROI covers the sensorimotor area, it should contain more information about the control of locomotor muscle synergy than the other ROIs. A previous study using transcranial magnetic stimulation and fMRI demonstrated that motor cortical regions activate muscle synergies of the leg and pelvic floor muscles, and that the regions have functional connectivity to widespread brain regions [43]. Based on these results and the results of the previous study, motor cortical regions with widespread cortical networks may be involved in the control of muscle synergies for leg muscles during walking, as well as the synergies of leg and pelvic floor muscles [43].

### Roles of slow cortical potentials in sensorimotor control

In the present study, slow cortical potentials in the delta bend (0.5–4 Hz) were used for our neural decoding method. Although such low-frequency cortical activity is associated with sleep [44], recent studies suggest that low-frequency cortical activity contains sensorimotor-related information. For example, delta band cortical activity plays a role in decision-making about somatosensory discrimination [45] and prediction of sensory events [46]. Additionally, neural decoding studies in humans have demonstrated that delta band activity is particularly informative for decoding kinematics parameters [27-32] and muscle activity [30]. In recent rodent studies, multisensory integration in widespread brain networks through slow cortical waves was suggested by calcium imaging [47]. As more direct evidence, a study on monkeys revealed intrinsic cyclic activity of slow cortical waves, functioning much like a spinal central pattern generator for locomotion, in the motor cortex and that slow waves synchronized upper-limb movements and muscle activity [48]. In addition, they demonstrated the slow cortical dynamics during sleep and under sedation. Given the task commonality between upper-limb movement and sleep, it is possible that the slow cortical dynamics are shared with walking. If the above-mentioned roles of slow cortical waves are conserved in humans, slow cortical waves may integrate muscle-synergy-related sensor information and be synchronized to muscle synergy activations. Therefore, locomotor muscle synergy activations could be decoded from slow waves in this study.

### Applicability to brain-machine interfaces

The decoding methodology and results of this study could contribute to the development of more effective locomotor rehabilitation approaches for patients with neural disorders. Recently, brain-machine interface (BMI) systems, which control stimulators that activate muscles through functional electrical stimulation (FES) based on cortical signals, have been used to aid recovery of movement in impaired patients [49]. As a new stimulation pattern of FES, muscle synergy-based stimulation patterns have been suggested for upper limb reaching [50] and locomotion [51]. The present results indicate that EEG signals contain information about the control of locomotor muscle synergies, providing fundamental information for effective neuroprosthetic systems based on a novel approach (e.g., BMI-FES with muscle synergy-based stimulation patterns) for restoring locomotion.

### Methodological considerations

Although EEG is a suitable method for examining brain activity during walking because of its high temporal resolution and mobility, the potential effects of movement artifacts should be considered. A recent study examined gait movement-related artifacts in EEG data by blocking the recording of electrophysiological signals (brain, eye, heart, and muscle activity) using a nonconductive layer (silicone swim cap) [52] and demonstrated that artifacts were smaller in electrodes in the central region (i.e., the vertex) compared with peripheral regions, because movement artifacts were caused by vertical head acceleration. In the present study, widely-distributed cortical activity, including that from central regions and peripheral regions (Figure 7A, 7B, and 7C), contributed to decoding. In addition, contribution to the decoding of central region ROI was larger than that of a peripheral ROI in a synergy type (Figure 7C). Thus, movement artifacts are not expected to have a major impact on the results.

Last, we used a primary components analysis (PCA)-based artifact rejection algorithm (ASR) to remove movement artifacts and other artifacts derived from muscle, heart, and eye activity. The ASR method removes high-variance artifact components from a dataset by comparison with a resting dataset [53]. This method has been utilized in studies recording EEG signals during walking, and its effectiveness has been confirmed in several studies [54, 55]. Therefore, the effects of movement artifacts on the current decoding results are not expected to be large.

## Conclusions

We demonstrated that low-frequency cortical waves are informative for the decoding of muscle synergy activity during walking, and that the decoding of individual muscle activity is based on muscle synergy-related cortical information. These results suggest that the cortex hierarchically controls locomotor muscle activity through muscle synergies. These novel findings advance our understanding of the neural control of human bipedal locomotion. Moreover, they demonstrate the feasibility of neural decoding of muscle synergy activation, supporting its future contribution to the development of effective brain-muscle neuroprostheses to restore walking in patients with mobility limitations.

## Acknowledgements

This work was supported by a Grant-in-Aid for Japan Society for the Promotion of Science Fellows (JSPS, #15J09583) to H.Y., Grant-in-Aid for Scientific Research (A) from JSPS to K.N. (#18H00818) and the Core Research for Evolutional Science and Technology (CREST) from Japan Science and Technology Agency (JST) to K.W. and K. N. (#JPMJCR14E4).

## Author Contributions

H.Y. and K.N. designed the experiment. H.Y. designed the current data analysis strategy. H.Y. and N.K. collected data. H.Y. performed analyses. H.Y., T.O., and K. N. drafted the manuscript. All authors interpreted the data, discussed the findings, and approved the final version of the manuscript

## Methods

### Experimental model and subject details Participants

Twelve healthy male volunteers (age, 23–31 years) participated in this study. Each participant provided written informed consent. The experiments were performed in accordance with the Declaration of Helsinki and with the approval of the Ethics Committee of the Graduate School of Arts and Sciences, University of Tokyo.

### Method details

#### Experimental design and setup

Participants walked on a treadmill (Bertec, Columbus, OH, USA) at 0.55 m/s for 7 min 30 seconds. The last seven minutes of data were used for the analysis. The slow walking speed was chosen based on two previous studies examining the effects of walking speed on movement artifacts in EEG signals [52, 54]: Kline et al. [52] used an experimental method to isolate and record independent movement artifacts with a silicone swim cap (nonconductive material), and reported large movement artifacts at walking speeds faster than 0.8 m/s. A study that analyzed relationships between head acceleration and motion artifacts in EEG signals indicated that recordings were robust at gait speeds below 3.0 km/h (0.83 m/s) [54]. As a static baseline condition, the participants sat on a chair for two minutes.

#### Data collection

Three-dimensional ground reaction forces (GRF) were recorded from force plates under the right and left belts of the treadmill (sampling rate: 1000 Hz). GRF data were smoothed with a low-pass filter (zero-lag Butterworth filter, 5 Hz cutoff). MATLAB 2016b (MathWorks, Natick, MA, USA) was used to perform all the post-processing analyses offline.

Surface electromyographic (EMG) signals were recorded from the following 13 leg muscles on the right side using a wireless EMG system (Trigno Wireless System, DelSys Inc., Boston, MA, USA): tensor fasciae latae (TFL), gluteus maximus (GM), gluteus medius (Gmed), sartorius (SART), biceps femoris (BF), semitendinosus (ST), rectus femoris (RF), vastus lateralis (VL), adductor magnus (AM), tibialis anterior (TA), peroneus longus (PL), soleus (SOL), and gastrocnemius medialis (MG). EMGs were amplified (with 300 gain preamplifier), band-pass filtered (20–450 Hz), and sampled at 1000 Hz.

A 64-channel EEG cap (Waveguard original, ANT Neuro b.v., Enschede, Netherlands) and a mobile EEG amplifier (eego sports, ANT Neuro b.v., Enschede, Netherlands) were used to record EEG signals at a sampling frequency of 500 Hz. Arrangement of the electrodes was according to the international 10–20 electrode system. EEG signals were referenced to CPz and a ground electrode was placed on AFz. Electrode impedances were kept below 30 kΩ (10 kΩ in most electrodes), which was substantially below the recommended impedance (below 50 kΩ) for the high-impedance EEG amplifier. Peripheral channels, which are prone to contamination by facial/cranial muscle activity and eye blinks, were removed from the offline analysis (channels labeled Fp, AF, FT, T, TP, O, PO, and F5-8, P5-8) [55], resulting in the 30 channels presented in Figure 1.

#### EMG processing and extraction of locomotor muscle synergies

Figure 2 shows an overview of our decoding methodology. From the recorded EMG signals, EMG envelopes and muscle synergies were used for the neural decoding analysis.

First, the recorded EMG data were high-pass filtered (zero-lag fourth-order Butterworth at 30 Hz), demeaned, full-wave rectified, and smoothed with a low-pass filter (zero-lag fourth-order Butterworth at 4 Hz cutoff) to obtain EMG envelopes [25]. EMG envelopes were resampled at 100 Hz. The amplitude of EMG envelopes for each muscle was normalized to the maximum value for that muscle during the walking task. Muscle synergies were extracted from the processed EMG envelopes using non-negative matrix factorization (NMF) [2, 3, 24-26]. For each participant, muscle synergies were extracted from the EMG dataset organized as a matrix with 13 muscles × 42000 variables (i.e., 100 Hz × 420 sec [7 min]). Using NMF, the EMG matrix (M) was decomposed into spatial muscle weightings (W), which correspond to the muscle synergies and their temporal activations (C) according to formula (1):

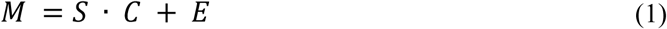

where *M* (*m* × *t* matrix, where *m* is the number of muscles and *t* is the number of samples in the EMG data matrix) is a linear combination of muscle synergies, *S* (*m* × Nsynergy matrix, where *N*_*synergy*_ is the number of muscle synergies), and their temporal activation patterns, *C* (*N*_*synergy*_ × *t* matrix), and *E* is the residual error matrix. The number of muscle synergies, *N*_*synergy*_, was determined by iterating each possible *N*_*synergy*_ from 1 to 10. For each *N*_*synergy*_, the goodness of fit was evaluated based on the variance accounted for (VAF) [56]. Based on the VAF, the optimal *N*_*synergy*_ was defined as the minimum value fulfilling two criteria: (1) the number of muscle synergies achieving VAF > 90% [56], and (2) the number to which adding an additional muscle synergy did not increase VAF by > 5% [57]. Then, we clustered the extracted muscle synergies using hierarchical clustering analysis to examine the extracted types of muscle synergies (Ward’s method, correlation distance) based on muscle weightings, as in our previous studies [3, 58, 59]. The gap statistic method was used to define the optimal number of clusters [60].

#### EEG pre-processing

In the current study, fluctuations in the amplitude of slow cortical potentials (0.5 – 4 Hz in the time domain) were used for the neural decoding analysis (Figure 2) based on a similar methodology used in previous studies [27-30, 32]. EEG data analysis was performed using custom programs in MATLAB incorporating functions of EEGLAB 14.1b [61]. The EEG signals were band-pass filtered between 0.5-100 Hz with a Butterworth filter (fourth-order). The “cleanline” function in EEGLAB was used to remove power line noise (50 Hz). Next, the EEG signals were resampled at 100 Hz. Then, we checked noisy EEG channels based on two criteria adopted from a previous study (Gwin et al., 2011): 1) standard deviation greater than 1000 μV, and 2) kurtosis of more than five standard deviations from the mean. In this study, no EEG electrode satisfied the criteria in all the participants. Since various types of artifacts were potentially introduced in the EEG data, we used an artifact rejection method called Artifact Subspace Reconstruction (ASR) [53] in EEGLAB to remove artifacts derived from walking, eye blinks, muscle, and heart activity. Next, the cleaned EEG signals were low-pass filtered at 4 Hz with a zero-phase Butterworth filter (fourth-order) and re-referenced to a common average reference. Finally, the amplitude of each electrode was normalized by calculating the standard z-score.

#### Neural decoding of muscle synergy and individual muscle activation

To continuously decode the activation of muscle synergies and individual muscles from the slow cortical potentials, we designed a time-embedded (10 lags, corresponding to 0 ms to −90 msec) linear decoding model, called the Wiener filter [28, 29, 62], for the muscle synergy and EMG envelope data. The linear model is given by:

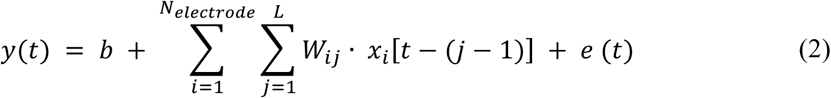

where *y*(*t*) is the predicted time series activation of each muscle synergy or EMG envelope at time *t, b* is the intercept, *N*_*electrode*_ (= 30) is the number of electrodes, *L* (=10) is the number of time lags, *x*(*t*) is the normalized slow cortical potentials at electrode *i* at time *t, W*_*ij*_ is the weights at electrode *i* and time lag *j*, and *e*(*t*) is the residual error. The parameters of the model were calculated with multidimensional generalized linear regression [28, 29, 62] using the “glm” function in MATLAB (Gaussian distribution condition). Neural decoders were designed separately for each participant and each decoded parameter (i.e., each muscle synergy and each EMG envelope).

For assessing the predictive accuracy of each decoder, a seven-fold cross-validation procedure was performed. Thus, the data recorded during the 7 min walking task were divided into 7 segments (1 min each). Six segments were used for training data while the remaining segment was used for testing the decoding model. This procedure was repeated for all possible combinations (i.e., seven times). Correlation coefficients (*r*) were calculated between the real activation and the decoded activation at each decoder in each iteration. To compare the overall decoding accuracy between the two types of decoders (muscle synergy decoders vs. individual muscle decoders), overall correlation values were calculated for each type per participant. To minimize the effects of skewness in the sampling distributions on the correlation coefficients, each correlation coefficient value was averaged after Fisher’s Z-transformation [63]. After averaging, the Z-values were back-transformed to the scale of Pearson’s r values.

Chance levels of neural decoding were evaluated by scrambling the EEG phase [64]. Phase-randomized EEG signals were generated by performing Fourier transform of a time series, and then the inverse Fourier transform was performed. The same decoding procedure was performed using the phase-randomized EEG signals. We generated 100 phase-randomized EEG datasets for each participant and performed neural decoding using each randomized dataset to obtain confidence intervals for the decoding accuracy.

#### Analysis of relationships between muscle synergy decoders and individual muscle decoders

We reconstructed individual muscle activations by summing the outputs of each decoded muscle synergy to test whether the variability in decoding accuracy in individual muscles would be reproduced by individual muscle activations indirectly decoded from muscle synergy activations decoded from muscle synergy decoders. The output of a decoded muscle synergy was explained by the product of the muscle weighting component and the decoded temporal activation pattern from the slow cortical potentials. Next, the decoding accuracy of the indirectly decoded individual muscle activation through the decoded muscle synergies were assessed.

To examine the weight of each muscle decoder (*W*_*muscle*_) based on those of the muscle synergy decoders (*W*_*syn*_), a 300-dimensional weight of an individual muscle decoder was reconstructed as a linear combination of the weights of the muscle synergy decoders with non-negative coefficients (*W*_*muscle*_*’*, conceptual schema presented in Figure 6C). The non-negative least squares problem was solved by the “lsqnonneg” function in MATLAB. The similarity of the weights of the original and reconstructed individual muscle decoders (i.e., *W*_*muscle*_ and *W*_*muscle*_*’*, respectively) was evaluated by Pearson’s r.

#### Contribution of each electrode to decoding

To evaluate the spatial contributions of cortical activity to predict muscle synergy activations, we calculated the contribution of each electrode from the weights of the decoding model as determined in a previous study [33]:

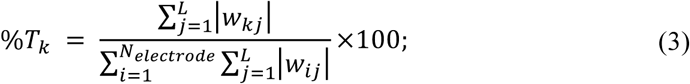

for all *k* from 1 to *N*_*electrode*_, where *%T*_*k*_ is the percentage contribution of each EEG electrode *k*.

In addition, we divided the electrodes into four major ROIs to examine the individual contribution of each area to the decoding. The ROIs were the frontal area (F3, F1, Fz, F2, F4, FC3, FC1, FCz, FC2, and FC4), central area (FC1, FCz, FC2, C3, C1, Cz, C2, C4, CP1, and CP2), lateral area (FC5, FC3, FC4, FC6, C5, C6, CP5, CP3, CP4, and CP6), and parietal area (CP3, CP1, CP2, CP4, P3, P1, Pz, P2, and P4). Using the same procedure as for the full electrodes, the decoding accuracy of each muscle synergy activation was separately calculated using the electrode set in each ROI.

#### Quantification and statistical analysis

The differences between the overall correlation values (i.e., decoding accuracy) between the two types of decoders (muscle synergy decoder vs. individual muscle decoder) were assessed using two-tailed paired t-tests. In addition, the differences in decoding accuracy between each ROI and the full electrode set were compared using repeated measures one-way analysis of variance (ANOVA) test with multiple t-tests with FDR correction for each muscle synergy type. For the statistical tests, the correlation values were transformed into Z-values using Fisher’s Z-transformation and the tests (i.e., t-test, ANOVA, multiple t-tests with FDR correction) were conducted on the Fisher’s Z-values. Statistical significance was set at *p* < 0.05.

